# Increased Nutrient Levels Enhance the Bacterial Contribution to an Algal-Bacterial Extracellular Matrix

**DOI:** 10.1101/2024.08.05.606509

**Authors:** Valeria Lipsman, Einat Segev

## Abstract

Microbial aggregation is central in environmental processes, such as marine snow and harmful marine mucilage events. Nutrient enrichment positively correlates with microbial aggregation. This correlation is largely attributed to the overgrowth of microalgae and the overproduction of agglomerating exopolysaccharides (EPS). However, recent studies highlight the significant contribution of bacterial EPS to algal-bacterial aggregation. Here, using controlled laboratory experiments and environmental metatranscriptomic analysis, we investigate the impact of nutrient enrichment on bacterial EPS production, while bacteria are in the context of their algal hosts. Our findings demonstrate a marked increase in bacterial EPS production in response to an increase of inorganic phosphorus and nitrogen levels, both in the lab and in the environment. These results highlight the interplay between nutrient regimes, bacterial physiology, and microbial aggregation in marine ecosystems and emphasize gaps in our understanding regarding the bacterial role in environmental processes that involve microbial aggregation.

## Introduction

Marine ecosystems are profoundly influenced by microbial communities, which play essential roles in nutrient cycling and primary production^1^. Microbes are key in the functioning of the biological carbon pump as they facilitate the transfer of carbon from the surface ocean to the deep sea^2^. Marine snow— comprised of organic and inorganic materials, including phytoplankton, fecal pellets, and microbial exopolymers—serves as a significant mediator of this carbon transport^3,4^. The microbial communities within marine snow are highly active, driving the decomposition and remineralization of organic matter^2^. Additionally, microbial aggregation processes can lead to harmful events such as marine mucilage, which involves the accumulation of gelatinous organic matter in coastal waters^5^. These mucilage events can adversely affect marine ecosystems by causing oxygen depletion, smothering benthic organisms, and disrupting aquaculture activities^5^. Understanding the role of microbial exopolysaccharides (EPS) in these aggregation processes is important for comprehending their impact on marine ecosystems^5^.

A positive correlation exists between elevated nutrient levels and enhanced microbial aggregation, both in terrestrial and marine ecosystems. The increased availability of nutrients appears to stimulate microbial growth, biofilm formation, and the subsequent aggregation of microorganisms, leading to the formation and stabilization of soil and marine aggregates^6–8^. In marine microbial communities, the synergistic effect of elevated temperature, pCO_2_, and nutrients has a significant impact on biofilm formation and community composition. This highlights the role of nutrient enrichment in enhancing microbial aggregation in marine environments, and indeed harmful mucilage events were found to develop under conditions of increased nutrient levels^9,10^.

Recently, we demonstrated that *Phaeobacter inhibens* bacteria contribute a specific EPS that promotes the formation of an algal-bacterial extracellular matrix (ECM) and enhances aggregation in co-culture with *Emiliania huxleyi* algae^11^. Building upon these findings, we sought to investigate whether elevated nutrient levels influence the bacterial contribution to EPS production. To address this question, we first examined the influence of elevated nutrient level on EPS production in controlled laboratory model systems. We simulated elevated nutrient regimes by manipulating the concentrations of inorganic phosphorus and nitrogen levels in the growth media of algal-bacterial co-cultures. By separately quantifying the bacterial and algal EPS, we observed a marked increase in both algal and bacterial EPS production in response to high levels of inorganic phosphorus and nitrogen. To gain insights into the ecological relevance of our observations, we conducted a metatranscriptomic analysis of natural bacterial communities that were sampled during an algal bloom induced by an influx of inorganic nutrients. Consistent with our laboratory results, the metatranscriptomic data revealed elevated expression of bacterial genes involved in EPS biosynthesis following the influx of inorganic nutrients. Furthermore, elevated expression of bacterial EPS genes correlated with the detection of various EPSs. These findings collectively support that bacteria exhibit increased EPS production following an influx of elevated nutrient levels, suggesting a bacterial contribution to the EPS pool under nutrient enriched conditions in marine environments. Our study highlights the interplay between nutrient levels, bacterial physiology, and microbial aggregation in marine ecosystems.

## Results

### Elevated levels of phosphate and nitrate increase algal and bacterial EPS production

We wished to assess the effect of inorganic phosphate (P) and nitrate (N) on EPS production of *P. inhibens* bacteria and *E. huxleyi* algae. Evaluating the impact of nutrient levels on bacteria and algae, separately, is challenging when cultivating co-cultures of this algal-bacterial pair. To address this challenge, we utilized specialized dual-cultivation chambers equipped with a 0.22 μm pore size membrane (Fig. 1a). This membrane separates algae and bacteria, allowing only small molecules to pass through. Consequently, the membrane maintains a shared chemical environment between the chambers while ensuring the separation of macromolecules that are produced by either algae or bacteria, such as EPS. Similar cultivation experiments have been previously conducted demonstrating that membrane-separated dual-cultivation chambers allow successful microbial metabolic exchange^12,13^. Algal and bacterial EPS production was tested under two P and N regimes. First, we added the recommended concentrations of nutrients to the L1-Si growth medium^14^, containing 36 μM P and 882 μM N (we denote this nutrient regime as 1×PN). These nutrient levels induce bloom-like growth of both algae and bacteria, resulting in cell densities of ∼10^6^ – 10^7^ cells/ml each^11^. Additionally, these P and N concentrations were previously shown to increase algal EPS production^15^. Therefore, this regime served as a control for nutrient concentrations that induce EPS production in *E. huxleyi*. In the second nutrient regime, we introduced lower nutrient concentrations (denoted as ¼×PN), containing 9 μM P and 220 μM N. These nutrient levels are similar to common oceanic conditions found in various marine environments^16^. At the early stationary phase of algal growth, cultures from each side of the dual-cultivation chambers were harvested. EPS production was assessed by the quantification of Alcian Blue (AB), a dye that binds to acidic EPS named Transparent Exopolymeric Particles (TEP), known to be produced by both algal and bacterial species^17^.

**Figure 1.**
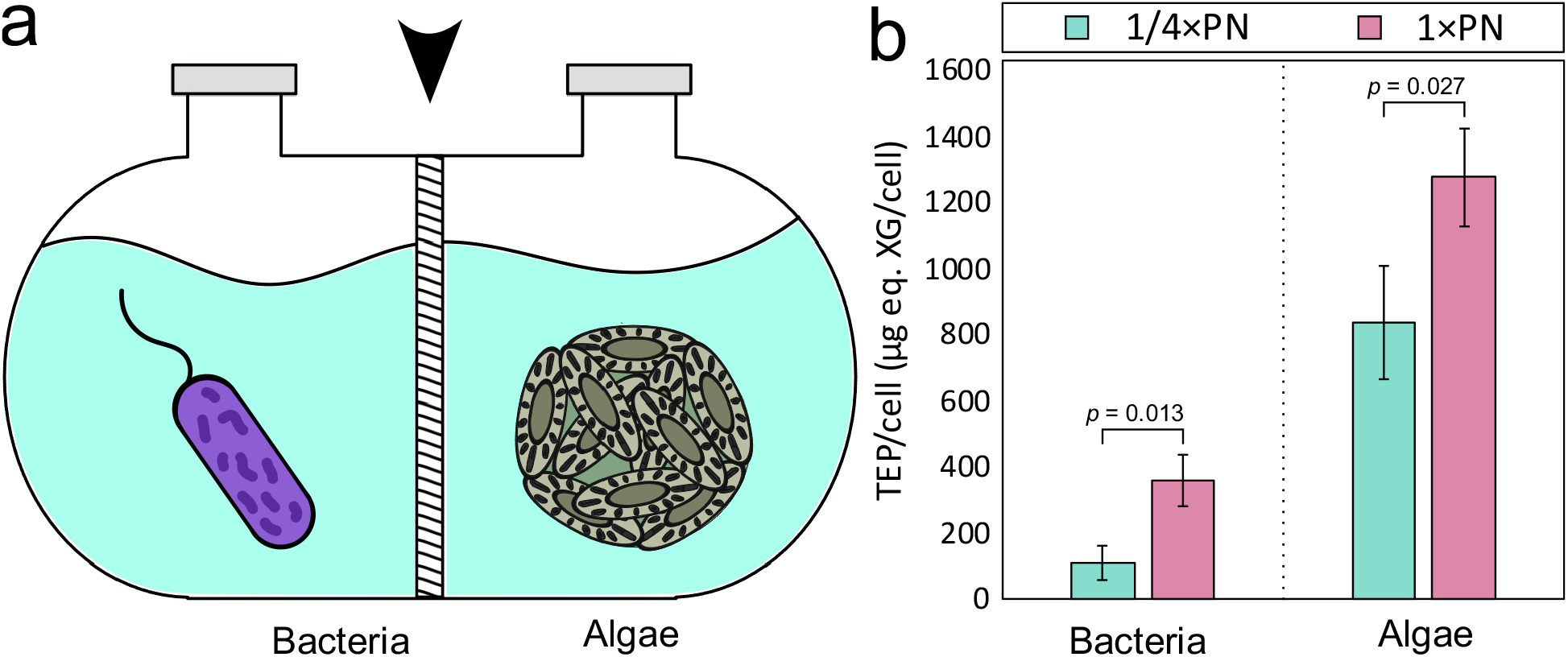
Elevated concentrations of phosphate and nitrate promote algal and bacterial EPS production. (a) A schematic representation of the dual-cultivation chambers, showing bacteria (left chamber) and algae (right chamber) separated by a membrane (marked by an arrowhead). (b) Bacterial (left) and algal (right) cultures were treated with ¼×PN (teal) or 1×PN (pink) nutrient regimes. EPS was quantified at early stationary phase (10 days) by measuring absorbance at 787 nm of AB bound to TEP and converting to Xanthan Gum (XG) equivalent weight. TEP/cell was calculated by dividing total TEP by the corresponding cell numbers (see Materials and Methods). Statistical significance was assessed using a one-tailed 2-sample t-test with three biological repeats, with p-values indicated above each organism.

Our data show that higher P and N concentrations affect both algal and bacterial EPS production (Fig. 1b). *E. huxleyi* TEP production was significantly higher (*p* = 0.027) under the 1×PN nutrient regime, compared to the ¼×PN regime, as previously demonstrated for other algal species^15^. Under both nutrient regimes, algae produce more TEP per cell than bacteria. However, bacteria also showed significantly higher (*p* = 0.013) TEP production under higher P and N concentrations, highlighting the potential bacterial contribution to aggregation at sea in response to elevated nutrient levels.

### Marine bacteria express EPS-related genes in response to a phosphate and nitrate influx

After observing increased bacterial EPS levels in response to elevated P and N concentrations in laboratory experiments, we aimed to examine if a similar bacterial response occurs in the environment. Recently, a study following the metatranscriptomics of bacterial populations during an algal bloom reported a peak of P and N environmental levels (Supplementary Fig. 1), prompting a secondary algal bloom with elevated levels of EPS (Supplementary Fig. 2)^18^. The increase in EPS was mainly attributed to algal production. We examined the publicly available metagenomics and metatranscriptomics data throughout the period of elevated EPS to assess whether upregulation of bacterial genes that are associated with EPS biosynthesis occurs after the peak of P and N.

From the collected metagenomic data, 251 representative metagenomic assembled genomes (MAGs) were previously generated and published^18^. Of these 251 MAGs, 7 MAGs belonged to Archaea and the rest were bacterial MAGs. To achieve a general view of the bacteria that are present in the analyzed samples, we constructed a phylogenetic tree of the MAGs (Fig. 2). The primary bacterial classes represented by the MAGs belonged to *Alphaproteobacteria, Gammaproteobacteria* and *Bacteroidia*. To assess the relative activity of the MAGs along the duration of the analysis, the transcriptomic data was mapped onto the representative MAGs and the counts were normalized by transcript per million (TPM). Relative activity was determined by mapping the relative TPM values to a certain MAG, and normalizing to the relative size of the MAG. During the sampling period, several MAGs exhibited higher activity during the second bloom that was induced by a nutrient influx in comparison to the initial natural bloom (Supplementary Figure 1). These active MAGs belonged to the *Alphaproteobacteria* and *Bacteroidia* classes.

**Figure 2.**
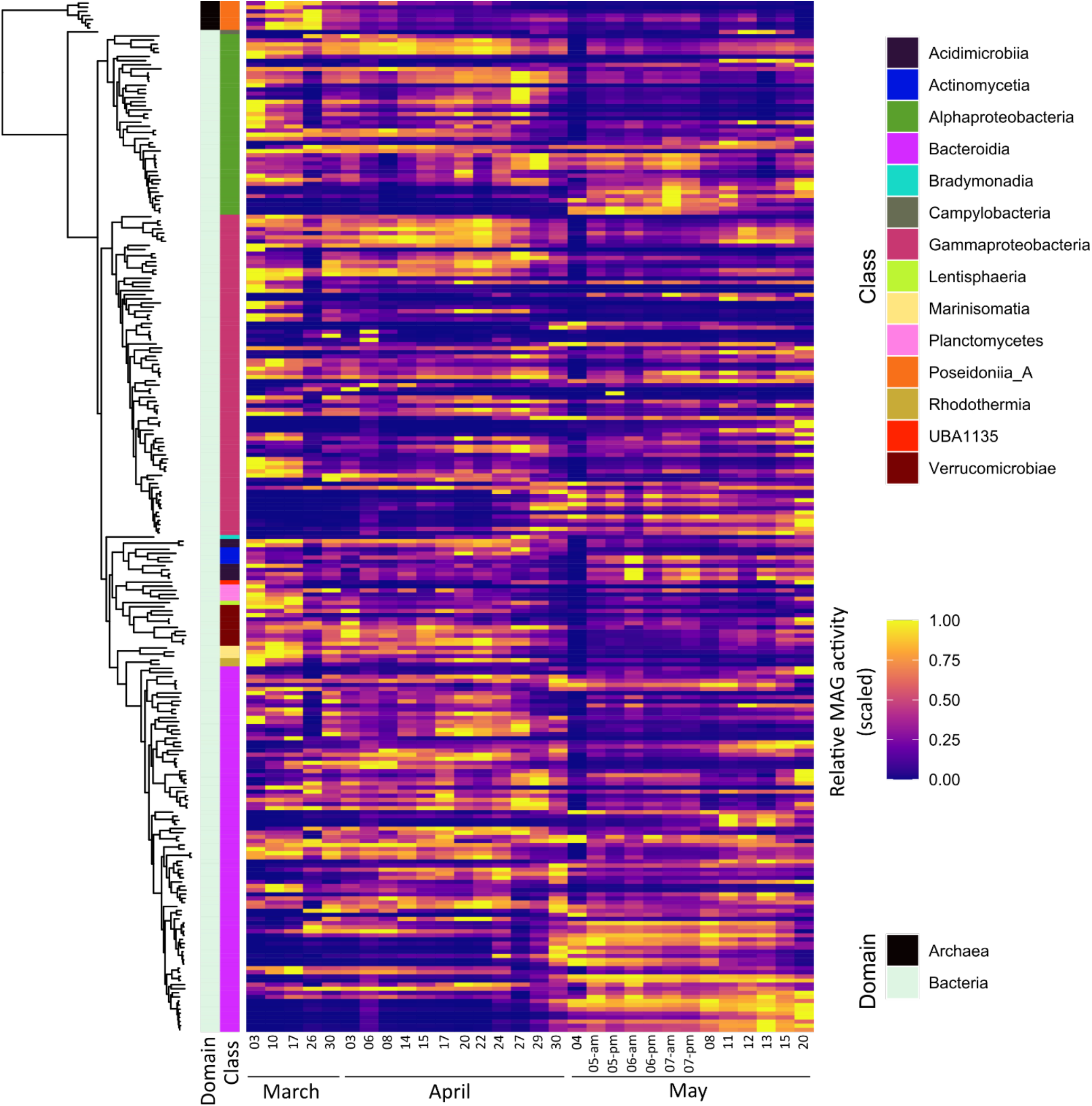
A peak of phosphate and nitrate increases the relative activity of specific bacterial groups in the environment. Phylogenetic tree of the 244 bacterial and 7 archeal representative MAGs (created by GTDB-Tk^19^) with domain and class affiliations (left). The relative activity (scaled) of each MAG is represented as a heatmap (right). Relative activity was calculated by summing the TPM values of all ORFs mapped to a MAG and dividing it by the sum of the TPM of all ORFs in a sample. This value was further normalized by the proportion of the MAG length compared to the total lengths of all MAGs. Data adapted from Sidhu, *et al*, 2023^18^.

To follow the expression of genes related to EPS production in the MAGs, we focused our analysis on bacterial genes involved in alginate, cellulose and wzx/wzy-dependent EPS biosynthesis^22^. These gene modules are involved in the production of many characterized bacterial EPSs, including acidic polysaccharides which potentially contribute to the TEP pool^23^. Additionally, we have previously shown that these bacterial modules are expressed by bacteria in co-culture with algae, contributing to algal-bacterial aggregation^11^. To characterize EPS biosynthesis modules in the representative MAGs, we searched for gene clusters of the key functions in this biosynthetic pathway (Supplemental table 1). We identified several MAGs that harbor gene clusters that putatively encode the majority of biosynthesis capacity of the different EPSs (Fig. 3). Several alginate and wzx/wzy-dependent gene modules exhibited higher expression during the second nutrient-induced algal bloom compared to the initial natural bloom phase. In contrast, the cellulose gene modules exhibited higher expression only at a single time point following the nutrient influx. Antibody-based measurements conducted by Sidhu *et al*. 2023^18^ demonstrated increased abundance of various EPSs from the 5^th^ to the 7^th^ of May following the nutrient influx (Supplemental Figure 2). Notably, on May 5^th^ and 6^th^, elevated expression was observed for genes belonging to all the three types of EPS modules-alginate, cellulose and wzx/wzy-dependent EPS. Given that bacteria are capable of producing diverse EPSs, with some resembling algal EPSs such as alginate and cellulose^22^, it cannot be ruled out that the EPSs that were previously identified are of both algal and bacterial origin. The co-occurrence of elevated expression of bacterial EPS modules and the increased EPS abundance suggests that bacteria may contribute to EPS production in response to P and N influx in the environment.

**Figure 3.**
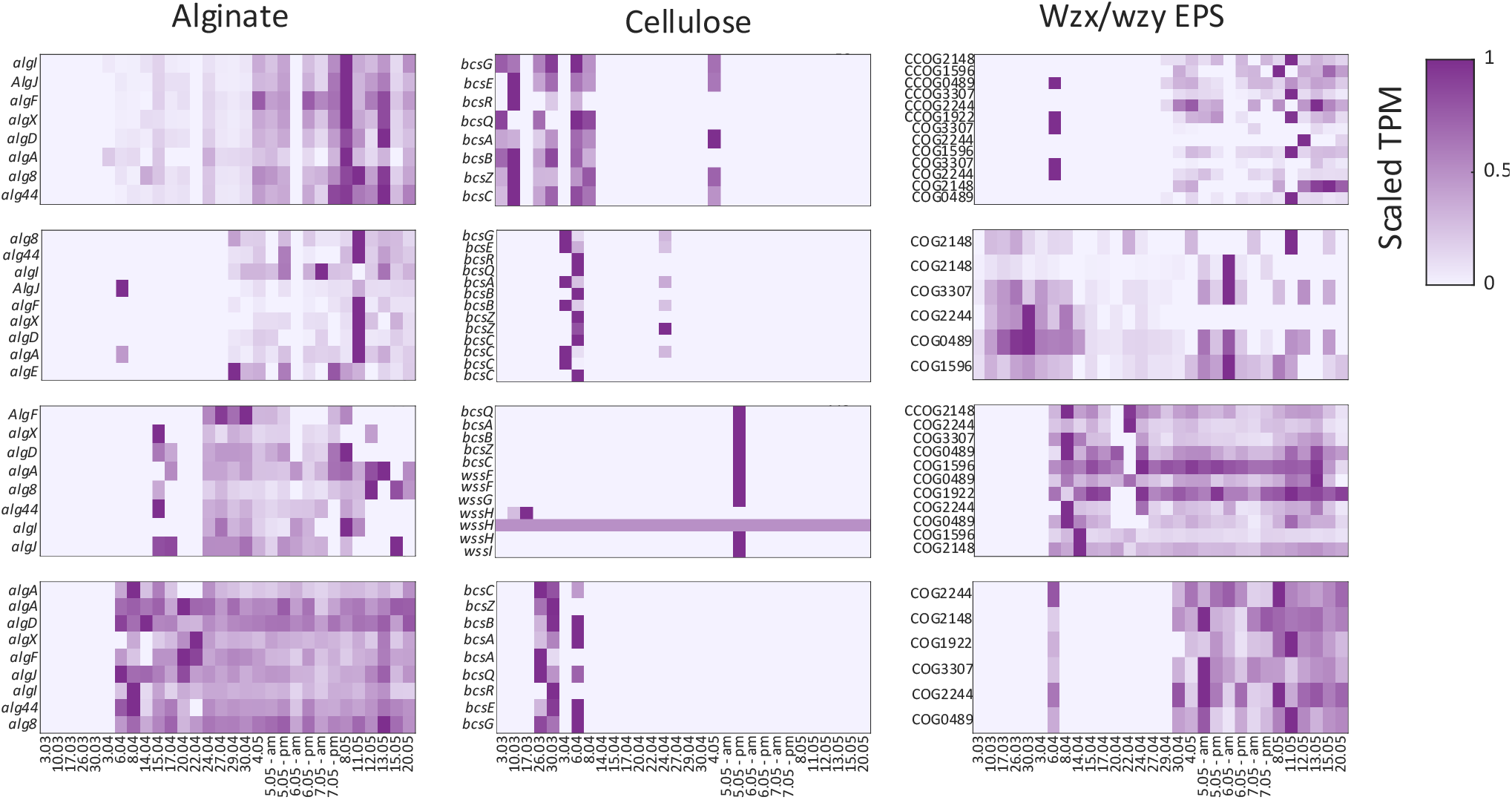
A peak of phosphate and nitrate increases the expression of bacterial genes related to EPS biosynthesis in the environment. The expression profiles of the gene clusters potentially involved in the biosynthesis of specific EPS in during the two algal bloom phases. Expression is presented as the scaled transcript abundance normalized by the abundance of 10 marker genes (MG) of each MAG. These values serve as the number of transcripts of a specific gene per MAG^20^ (see Materials and Methods). Gene clusters related to the biosynthesis of alginate, cellulose and wzx/wzy dependent EPS are shown for different MAGs. Shown are the genes (for alginate and cellulose) or the OG numbers associated with the wzx/wzy depended EPS biosynthesis genes (annotated using EggNOG-mapper v2^21^). Data adapted from Sidhu, *et al*, 2023^18^.

## Discussion

In this study, we demonstrate that bacterial EPS production increases following a nutrient influx in both laboratory settings and marine environments. Our results demonstrate that higher concentrations of P and N lead to increased EPS production in both *E. huxleyi* algae and *P. inhibens* bacteria. Specifically, *E. huxleyi* algae showed a significant increase in TEP production under the higher nutrient regime compared to the lower nutrient regime. This observation aligns with previous studies that demonstrated nutrient-induced EPS production in algal species^15^. Notably, while algae produced more TEP per cell than bacteria, bacteria also showed a significant increase in TEP production under elevated nutrient conditions. Of note, the interaction between algae and heterotrophic bacteria plays a significant role in shaping microbial communities. Algae release molecules that influence bacterial physiology, and heterotrophic bacteria depend on algae to release exudates and transform inorganic compounds like nitrite into utilizable forms^24^. Therefore, elevated nutrient levels may have a direct impact on bacterial EPS metabolism, or an indirect effect through the algal host.

Our metagenomic and metatranscriptomic analyses of a marine algal bloom revealed that several bacterial MAGs exhibited higher activity during the secondary nutrient-induced bloom phase compared to the initial natural bloom. The previously published data that was analyzed here showed that the composition and abundance of both algal and bacterial species varied significantly between the two algal bloom phases^18^. It was previously suggested that the availability of algal polysaccharides can impact the structure of bacterial communities. The algal polysaccharides provide essential nutrients for heterotrophic bacteria, and can form a dense matrix that facilitates bacterial attachment and aggregation^25^. In turn, bacteria in proximity to their algal hosts can contribute to the EPS pool, promoting further algal-bacterial agglomeration^11^. The elevated expression of bacterial EPS modules in environmental samples suggests a potential bacterial contribution to marine aggregation processes specifically in nutrient-rich environments. Our understanding of algal-bacterial aggregation in the environment is partial at best. The current study highlights the complex dynamics between algal and bacterial communities and emphasizes the potential impact of nutrient influxes on the ecology of microbial populations.

Influxes of waters that carry enriched N and P levels can impact the composition and spatial traits of various aquatic microbial populations^22,23^. Eutrophication, the increase in nutrient levels, was reported to encourage growth of harmful species, change trophic networks, and alter the diversity and richness of the ecosystem^25^. Elevated levels of phosphate and nitrogen in marine environments are known to play a significant role in promoting marine mucilage events^26^. Marine mucilage can carry distinct bacterial communities^5^ which can play a role in these events by producing EPS and further contributing to mucilage formation. These mucus layers can have profound ecological impacts, including the disruption of marine food webs, the suffocation of marine organisms, and the alteration of biogeochemical cycles^5^. Understanding the bacterial role in these events is crucial for developing strategies to manage and mitigate their occurrence in marine ecosystems.

## Materials and Methods

### Microbial strains and growth conditions

The bacterial strain *Phaeobacter inhibens* DSM 17395 was acquired from the German Collection of Microorganisms and Cell Cultures (DSMZ, Braunschweig, Germany). Bacteria were cultivated by inoculation from a frozen stock (−80°C) onto ½ YTSS agar plates containing 2 g yeast extract, 1.25 g tryptone, and 20 g sea salt per liter (all obtained from Sigma-Aldrich, St. Louis, MO, USA). Plates were incubated at 30 °C for 2 days. Individual colonies were used to initiate bacterial cultures in artificial seawater (ASW) medium. Bacterial starters were supplemented with CNPS to support bacterial growth. CNPS consisted of L1-Si medium (detailed below)^24,27^ supplemented with glucose (5.5 mM), Na_2_SO_4_ (33 mM), NH_4_Cl (5 mM), and KH_2_PO_4_ (2 mM), all sourced from Sigma-Aldrich. The cultures were incubated at 30 °C with continuous shaking at 130 rpm for 2 days until the start of the experiments (day 0).

The axenic algal strain *Emiliania huxleyi* CCMP3266 was obtained from the National Center for Marine Algae and Microbiota (Bigelow Laboratory for Ocean Sciences, Maine, USA). Algae were cultivated in L1 medium based on the protocol by Guillard and Hargraves^14^. Notably, Na_2_SiO_3_ was omitted as per cultivation recommendations for this algal strain, and the medium was designated as L1-Si. Algal cultures were maintained in standing conditions within a growth room at a temperature of 18 °C under a light/dark cycle of 16/8 h. Light intensity during the illuminated period was maintained at 150 mmoles/m^2^/s. The absence of bacteria in axenic algal cultures was periodically verified through plating on ½ YTSS plates and microscopic observations.

### Microbial growth in dual-cultivation chambers

Dual-cultivation chambers were custom made based on Vallet, *et al*, 2019^13^ (Bein M.LTD, Tel Aviv, Israel). A 100 mm 0.22 μm GV Durapore membrane filter (from Sigma-Aldrich) divided between chambers in the dual-cultivation setup. Each side of the split chamber contained 30 ml of medium with the regular concentrations of vitamins and trace metals of L1-Si medium. Phosphate (NaH_2_PO_4_· H_2_O, denoted as P) and nitrate (NaNO_3_, denoted as N), were either supplied in ¼ of the regular L1-Si (denoted as ¼×PN), or regular L1-Si concentrations (denoted as 1×PN).

#### Algal inoculation

Algal cell concentrations from a late exponential phase culture were enumerated using a CellStream CS-100496 flow cytometer (Merck, Darmstadt, Germany) with 561 nm excitation and 702 nm emission. Each sample involved recording 50,000 events. An inoculum of 10^4^ algal cells was introduced into one side of the split chamber.

#### Bacterial inoculation

*P. inhibens* bacteria (prepared as previously described) were quantified by measuring OD_600_ values using an Ultrospec 2100 pro spectrophotometer (Biochrom, Cambridge, UK) with plastic cuvettes. Cell numbers were adjusted based on a pre-existing calibration curve to inoculate a concentration of 10^2^ cells/ml in 30 ml of medium, resulting in 10–100 colony forming units (CFU) per mL, which were verified by plating serial dilutions (in ASW) on ½ YTSS plates. Bacteria were inoculated into one side of the split chambers following four days of algal growth in the other side. Following bacterial inoculation, the split chambers were incubated for 10 additional days under the same conditions as described above for algal cultivation.

#### Monitoring bacterial and algal growth

Bacterial concentrations in split chambers were determined by counting CFU/mL. The cultures were serially diluted in ASW and plated on ½ YTSS plates. The plates were incubated at 30 °C for 2 days, and subsequently, colonies were enumerated to calculate bacterial concentration in CFU/mL. Algal cell concentrations were quantified using a flow cytometer, as previously described. Cultures were thoroughly mixed and subsequently, 100 μL of the samples were transferred to round-bottom 96-well plates. Each sample was recorded for 1 minute to capture a total of > 50,000 events. Cross-contamination was examined by monitoring the CFU/mL and flow cytometry samples of both algal and bacterial sides at the time of initial inoculation and sample collection.

### TEP measurement

Alcian Blue (AB) staining was performed following published protocols^17^ using a staining solution with 200 mg L^-1^ of Alcian Blue (8GX, Sigma-Aldrich). The solution was prepared in acidified ultrapure water, adjusting the pH to 2.5 with glacial acetic acid. Each culture was filtered on Isopore™ 25 mm 0.4 μm PC membrane (Sigma-Aldrich) in three biological repeats, with 6 mL and 2 mL of culture for bacteria and algae, respectively. Following filtration, the membranes were washed twice with 1 mL of DDW and stained with 1 mL AB for 10 seconds. The dye was removed, and the filters were washed twice with 1 mL of DDW. Xanthan Gum (XG, Sigma-Aldrich) was used for calibration by dissolving 75 mg XG in 1 L DDW, mixing for 3 hours. Serial dilutions where prepared by mixing 0.125, 0.250, 0.500, 0.750, and 1 mL of the XG with DDW to a final volume of 1 mL. The dilutions were then mixed with 0.5 mL AB, filtered, and washed as described above. The stained filters were transferred to 15 mL falcon tubes with 80% sulfuric acid and mixed for 2 hours. Absorbance was measured at 787 nm wavelength and TEP_0.4μm_ concentrations were calculated as follows:

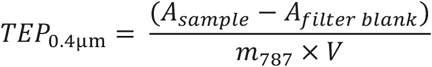

Where TEP_0.4μm_ is expressed as XG μg/μL. m787 is the slope of the calibration curve of XG. V is the volume of sample filtered. A_sample_ is the absorbance of AB extracted from a sample filter. A_filter blank_ is the absorbance of AB extracted from a filter used with only medium. As blank, 80% sulfuric acid was used. Each result was divided by the corresponding Cells/μL or CFU/μL values to calculate TEP_0.4μm_/Cell values.

### MAG analysis

A collection of 251 MAGs was previously assembled by Sidhu, *et al*, 2023^18^, and re-analyzed in the current study. MAG phylogenetic affiliations were re-assigned using GTDB-Tk v2.3.2^19^ with GTDB v214. A phylogenetic tree was constructed using de_novo_wf command of GTDB-Tk v2.3.2^19^. The tree was extracted and visualized using the ape and ggtree packages in R v4.3.1. ORFs prediction was conducted using Prodigal^28^. The ORFs were annotated using EggNOG-mapper v2^21^. HMM homology was searched with HMMER3^29^ against the Pfam-A HMM database^30^. HMM models of biosynthetic genes related to EPS biosynthesis were acquired from the KEGG KofamKOALA^31^ and EggNOG v5.0^32^ databases. The hmm models were used to search for orthologos in the representatives MAGs with HMMER3^29^. Manual annotation of the biosynthetic EPS modules was conducted by searching gene clusters that contain the ORFs associated with the key genes of the core functions of the biosynthetic pathways^22^, summarized in Supplementary Table 1.

## Metatranscriptomics analysis

Raw sequence data was acquired for 30 time points from the European Nucleotide Archive (accession PRJEB52999)^18^. Ribosomal RNA reads were removed using SortMeRNA v4.3.6^33^ searched against the smr_v4.3_default_db database. Messenger RNA reads were quality trimmed and end repaired using the bbduk and repair scripts from the BBMap v39.01 (https://sourceforge.net/projects/bbmap/). Reads with a minimum length of 70 bp were then mapped to all 251 representative MAGs using Bowtie2 v2.5.1^34^. Counts were calculated by FeatureCounts^35^. First, a transcript per million (TPM) normalization was conducted for all samples as follows^36^:

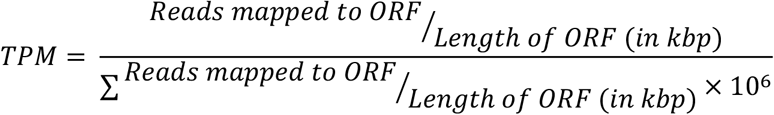

To assess MAG activity, the sums of TPM values of each MAG in a given sample were normalized as follows^37^:

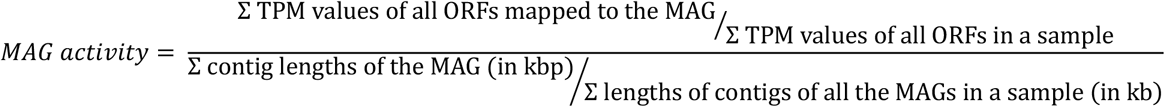

Top 15 relatively active MAGs were selected for further analysis. For expression analysis of polysaccharide -related genes in a MAG, the counts were first transformed with marker genes (MG) normalization, as described previously^20^, using 10 universal single-copy phylogenetic MGs as OGs^38,39^: COG0012, COG0016, COG0018, COG0172, COG0215, COG0495, COG0525, COG0533, COG0541, and COG0552. Genes were extracted using FetchMGs v1.2 (available at http://motu-tool.org/fetchMG.html). The normalization was conducted as follows:

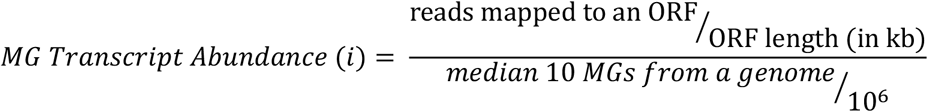

As the MGs were shown to be constitutively expressed across many different conditions^39^, MGs normalized transcript abundance can be interpreted as the relative per-MAG number of transcripts of a given ORF^20^. MAGs containing clusters of genes related to core functions, which are highly conserved, were selected and transcript abundance was plotted with MATLAB R2023b.

## Acknowledgments

We thank all members of the Segev lab for insightful comments and discussions. This study was supported by funds received from the European Research Council (ERC StG 101075514) and the de Botton Center for Marine Science, granted to E.S.

## Competing interests

The authors declare no competing interests.

## Authors Contributions

V.L. and E.S. designed the study. V.L. performed experiments. V.L. and E.S. analyzed results and wrote the manuscript.

## Supplementary Data

**Supplementary Table 1.** The core functions of the genes in biosynthesis of different EPSs, with KEGG, OG and main Pfam ID domains

| EPS | Gene | KEGG | OG | PFAM |
| --- | --- | --- | --- | --- |
| Alginate | alg8 | K19290 | COG1215 |  |
|  | alg44 | K19291 |  | PilZ |
|  | algK | K19292 | COG0790 | Sel1 |
|  | algX | K19293 |  | ALGX |
|  | algI | K19294 | COG1696 | MBOAT |
|  | algJ | K19295 |  | ALGX |
|  | algF | K19296 |  | AlgF |
|  | algE | K16081 |  | Alginate_exp |
|  | algG | K01795 | COG3420 | Beta_helix |
|  | algL | K01729 |  | Alginate_lyase |
| Cellulose | bcsA | K00694 | COG1215 | Cellulose_synt |
|  | bcsB | K20541 | COG1215 | BcsB |
|  | bcsZ | K20542 | COG3405 | Glyco_hydro_8 |
|  | bcsC | K20543 | COG5010 | BCSC_C |
|  | bcsG |  | COG2194 | CBP_BcsG |
|  | bcsE |  |  | CBP_BcsE |
|  | bcsR |  |  | CBP_BcsR |
|  | bcsQ |  |  | CBP_BcsQ |
| Wzx/Wzy | Primary glycosyl transferase |  | COG2148 | Bac_transf |
|  | Repeating unit flippase |  | COG2244 | Polysacc_synt |
|  | Repeating unit polymerase |  | COG3307 | Wzy_C |
|  | Polysaccharide co-polymerase |  | COG0489 | Wzz |
|  | Outer membrane polysaccharide export |  | COG1596 | Poly_export |
|  | Repeating unit tranferase |  | COG1922 | Glyco_tran_WecG |

**Supplementary Figure 1.**
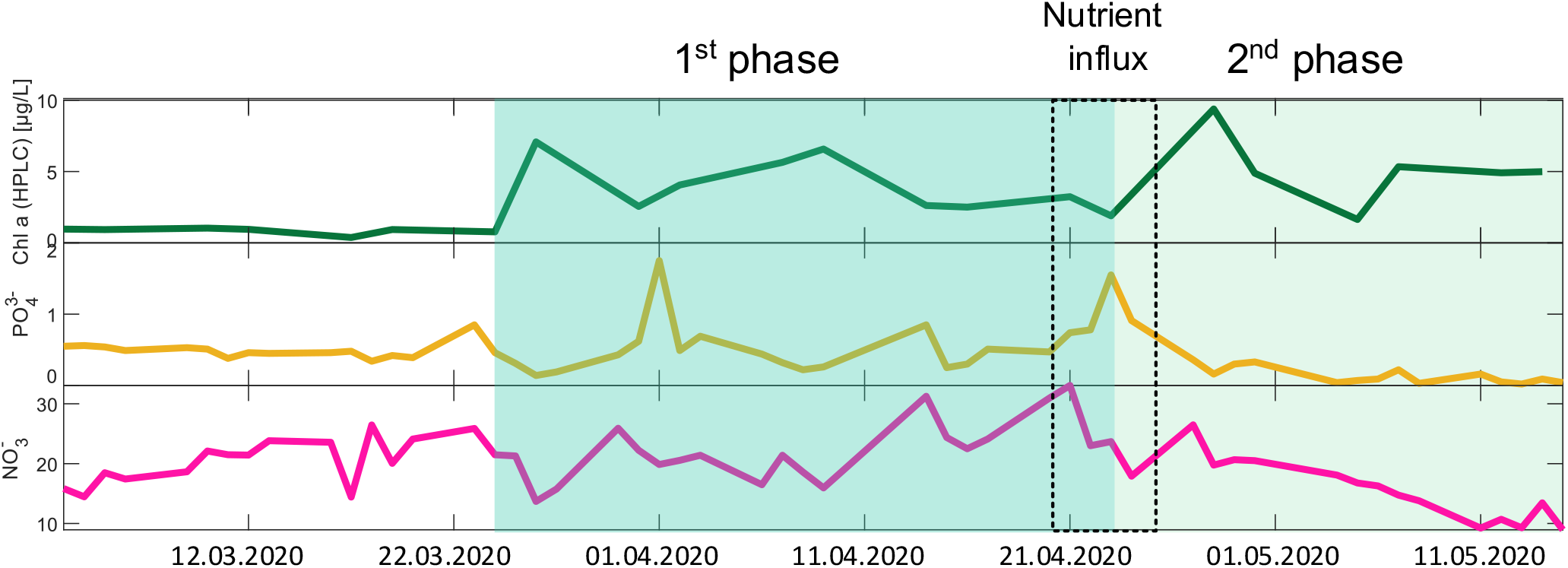
Environmental data during the sampling period. Measurements of Chlorophyll *a* (top), NO_3_^-^ (bottom) and PO_4_^3-^ (middle) concentrations during the period of the analysis. Marked are the two algal bloom phases and the time of the nutrient influx. Data adapted from Sidhu, *et. al*., 2023^18^.

**Supplementary Figure 2.**
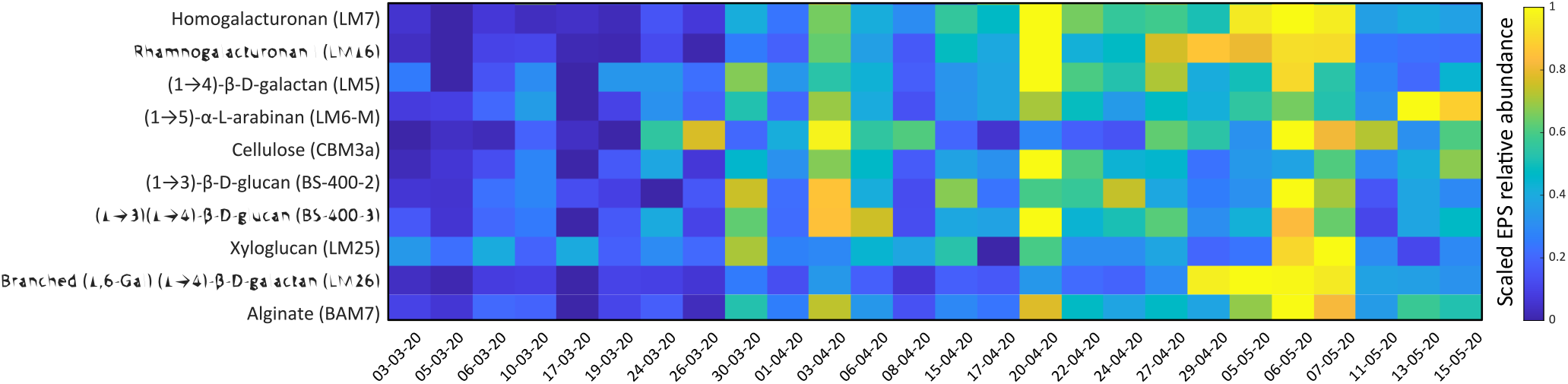
Abundance of specific EPSs during the sampling period. Antibody-based measurements of different dissolved EPSs during the period of the analysis. Shown are various EPS and the monoclonal antibody used to target the EPS. EPS was extracted from the high molecular weight dissolved organic matter (HMWDOM) of each sample, using EDTA as solvent. Values indicate antibody signal intensity. Color bar shows values that were scaled for each EPS and are presented as the fraction of the highest measured abundance. Data adapted from Sidhu, *et. al*., 2023^18^.

## Notes

### Competing Interest Statement

The authors have declared no competing interest.

## References

1. Field, C. B., Behrenfeld, M. J., Randerson, J. T. & Falkowski, P. Primary production of the biosphere: Integrating terrestrial and oceanic components. Science (80-.). 281, 237–240 (1998).

2. Ploug, H. & Grossart, H. P. Bacterial growth and grazing on diatom aggregates: Respiratory carbon turnover as a function of aggregate size and sinking velocity. Limnol. Oceanogr. 45, 1467–1475 (2000).

3. Turner, J. T. Zooplankton fecal pellets, marine snow, phytodetritus and the ocean’s biological pump. Progress in Oceanography vol. 130 205–248 (2015).

4. Simon, M., Grossart, H. P., Schweitzer, B. & Ploug, H. Microbial ecology of organic aggregates in aquatic ecosystems. Aquatic Microbial Ecology vol. 28 175–211 (2002).

5. Danovaro, R., Fonda Umani, S. & Pusceddu, A. Climate Change and the Potential Spreading of Marine Mucilage and Microbial Pathogens in the Mediterranean Sea. PLoS One 4, 1–8 (2009).

6. Wilpiszeski, R. L. et al. Soil Aggregate Microbial Communities: Towards Understanding Microbiome Interactions at Biologically Relevant Scales. Applied and Environmental Microbiology vol. 85 (2019).

7. Baragi, L. V. & Anil, A. C. Synergistic effect of elevated temperature, pCO2 and nutrients on marine biofilm. Mar. Pollut. Bull. 105, 102–109 (2016).

8. Chen, J. et al. Soil Aggregation Shaped the Distribution and Interaction of Bacterial-Fungal Community Based on a 38-Year Fertilization Experiment in China. Front. Microbiol. 13, (2022).

9. Savun-hekimoglu, B. & Gazioğlu, C. Mucilage Problem in the Semi-Enclosed Seas: Recent Outbreak in the Sea of Marmara. Int. J. Environ. Geoinformatics 8, 402–413 (2021).

10. Malone, T. C. & Newton, A. The Globalization of Cultural Eutrophication in the Coastal Ocean: Causes and Consequences. Frontiers in Marine Science vol. 7 (2020).

11. Lipsman, V., Shlakhter, O., Rocha, J. & Segev, E. Bacteria contribute exopolysaccharides to an algal-bacterial joint extracellular matrix. npj Biofilms Microbiomes 10, (2024).

12. Thøgersen, M. S., Melchiorsen, J., Ingham, C. & Gram, L. A Novel Microbial Culture Chamber Co-cultivation System to Study Algal-Bacteria Interactions Using Emiliania huxleyi and Phaeobacter inhibens as Model Organisms. Front. Microbiol. 9, 361675 (2018).

13. Vallet, M. et al. The oomycete Lagenisma coscinodisci hijacks host alkaloid synthesis during infection of a marine diatom. Nat. Commun. 10, (2019).

14. Guillard, R. R. L. & Hargraves, P. E. Stichochrysis immobilis is a diatom, not a chrysophyte. Phycologia 32, 234–236 (1993).

15. Buzzelli, E., Gianna, R., Marchiori, E. & Bruno, M. Influence of nutrient factors on production of mucilage by Amphora coffeaeformis var. perpusilla. Cont. Shelf Res. 17, 1171–1180 (1997).

16. Raven, J. A., Gobler, C. J. & Hansen, P. J. Dynamic CO2 and pH levels in coastal, estuarine, and inland waters: Theoretical and observed effects on harmful algal blooms. Harmful Algae 91, (2020).

17. Passow, U. Transparent exopolymer particles (TEP) in aquatic environments. Progress in Oceanography vol. 55 287–333 (2002).

18. Sidhu, C. et al. Dissolved storage glycans shaped the community composition of abundant bacterioplankton clades during a North Sea spring phytoplankton bloom. Microbiome 11, (2023).

19. Chaumeil, P. A., Mussig, A. J., Hugenholtz, P. & Parks, D. H. GTDB-Tk: A toolkit to classify genomes with the genome taxonomy database. Bioinformatics 36, 1925–1927 (2020).

20. Salazar, G. et al. Gene Expression Changes and Community Turnover Differentially Shape the Global Ocean Metatranscriptome. Cell 179, 1068-1083.e21 (2019).

21. Cantalapiedra, C. P., Herņandez-Plaza, A., Letunic, I., Bork, P. & Huerta-Cepas, J. EggNOG-mapper v2: Functional Annotation, Orthology Assignments, and Domain Prediction at the Metagenomic Scale. Mol. Biol. Evol. 38, 5825–5829 (2021).

22. Vandana, G. & Das, S. Genetic regulation, biosynthesis and applications of extracellular polysaccharides of the biofilm matrix of bacteria. Carbohydr. Polym. 291, 119536 (2022).

23. Schmid, J., Sieber, V. & Rehm, B. Bacterial exopolysaccharides: Biosynthesis pathways and engineering strategies. Front. Microbiol. 6, (2015).

24. Segev, E. et al. Dynamic metabolic exchange governs a marine algal-bacterial interaction. Elife 5, (2016).

25. Bar-Zeev, E., Berman-Frank, I., Girshevitz, O. & Berman, T. Revised paradigm of aquatic biofilm formation facilitated by microgel transparent exopolymer particles. Proc. Natl. Acad. Sci. U. S. A. 109, 9119–9124 (2012).

26. Volkan Oral, H. Environmental statistical analysis on the impacts of marine mucilage on some seawater quality parameters. doi:10.30897/ijegeo.1187859.

27. Zech, H. et al. Growth phase-dependent global protein and metabolite profiles of Phaeobacter gallaeciensis strain DSM 17395, a member of the marine Roseobacter-clade. Proteomics 9, 3677–3697 (2009).

28. Hyatt, D. et al. Prodigal: Prokaryotic gene recognition and translation initiation site identification. BMC Bioinformatics 11, (2010).

29. Eddy, S. R. A new generation of homology search tools based on probabilistic inference. Genome Inform. 23, 205–211 (2009).

30. Finn, R. D. et al. The Pfam protein families database: Towards a more sustainable future. Nucleic Acids Res. 44, D279–D285 (2016).

31. Aramaki, T. et al. KofamKOALA: KEGG Ortholog assignment based on profile HMM and adaptive score threshold. Bioinformatics 36, 2251–2252 (2020).

32. Huerta-Cepas, J. et al. EggNOG 5.0: A hierarchical, functionally and phylogenetically annotated orthology resource based on 5090 organisms and 2502 viruses. Nucleic Acids Res. 47, D309–D314 (2019).

33. Kopylova, E., Noé, L. & Touzet, H. SortMeRNA: Fast and accurate filtering of ribosomal RNAs in metatranscriptomic data. Bioinformatics 28, 3211–3217 (2012).

34. Langmead, B. & Salzberg, S. L. Fast gapped-read alignment with Bowtie 2. Nat. Methods 9, 357–359 (2012).

35. Liao, Y., Smyth, G. K. & Shi, W. FeatureCounts: An efficient general purpose program for assigning sequence reads to genomic features. Bioinformatics 30, 923–930 (2014).

36. Wagner, G. P., Kin, K. & Lynch, V. J. Measurement of mRNA abundance using RNA-seq data: RPKM measure is inconsistent among samples. Theory Biosci. 131, 281–285 (2012).

37. Mickol, R. L. et al. Metagenomic and metatranscriptomic characterization of a microbial community that catalyzes both energy-generating and energy-storing electrode reactions. Appl. Environ. Microbiol. 87, (2021).

38. Sunagawa, S. et al. Metagenomic species profiling using universal phylogenetic marker genes. Nat. Methods 2013 1012 10, 1196–1199 (2013).

39. Milanese, A. et al. Microbial abundance, activity and population genomic profiling with mOTUs2. Nat. Commun. 10, (2019).

