## Supplemental figure S1 and S2 for "Increased Nutrient Levels Enhance the Bacterial Contribution to an Algal-Bacterial Extracellular Matrix"

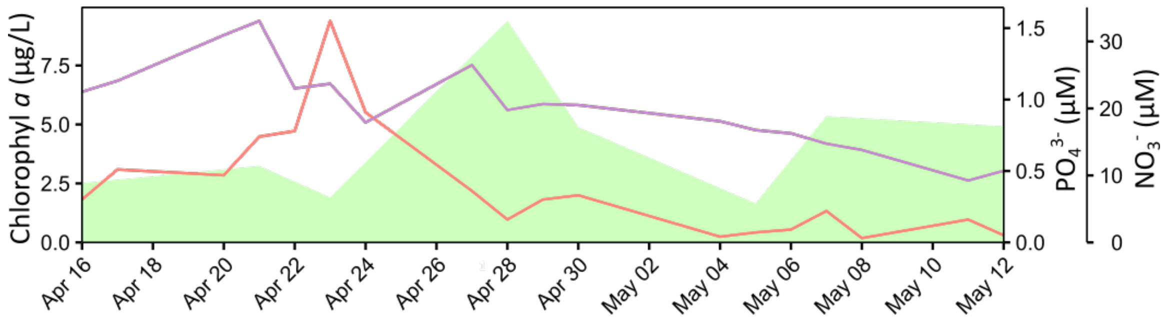


**Supplementary Figure 1. Environmental data during the sampling period.** Measurements of Chlorophyll *a* (green area), $NO_3^-$ (purple line) and $PO_4^{3-}$ (orange line) concentrations during the period of the analysis. Data adapted from Sidhu, *et. al.*, 2023[1].

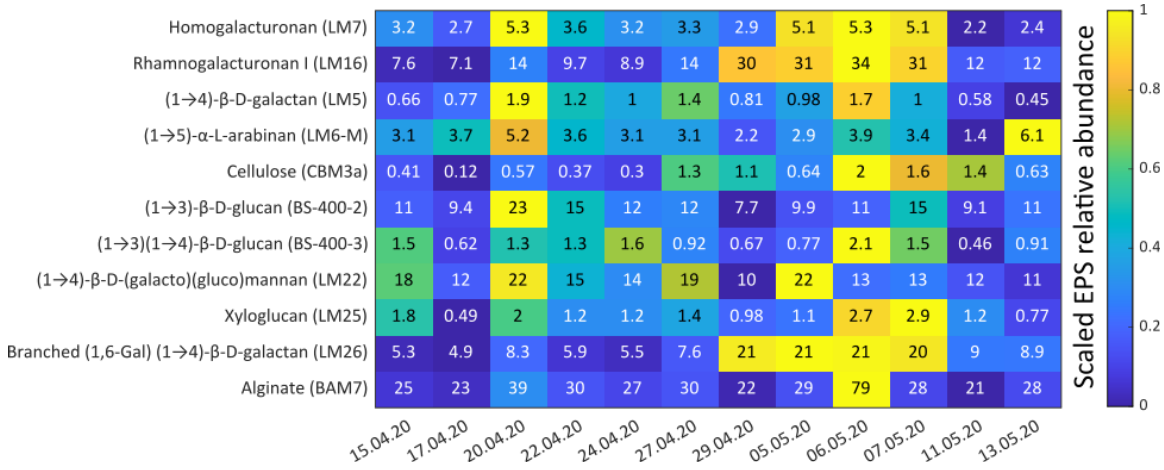


**Supplementary Figure 2. Abundance of specific EPSs during the sampling period.** Antibody-based measurements of different dissolved EPSs during the period of the analysis. Shown are various EPS and the monoclonal antibody used to target the EPS. EPS was extracted from the high molecular weight dissolved organic matter (HMWDOM) of each sample, using EDTA as solvent. Values indicate antibody signal intensity. Color bar shows values that were scaled for each EPS and are presented as the fraction of the highest measured abundance. Data adapted from Sidhu, *et. al.*, 2023[1].

### Supplementary References

1. Sidhu, C. *et al.* Dissolved storage glycans shaped the community composition of abundant bacterioplankton clades during a North Sea spring phytoplankton bloom. *Microbiome* **11**, (2023).
